# Structural profiling of the pneumolysin epitope landscape uncovers a cross-species neutralising site across cholesterol-dependent cytolysins

**DOI:** 10.64898/2026.02.27.708574

**Authors:** Di Tang, Indrė Kučinskaitė-Kodzė, Joel Ströbaek, Carlos Gueto-Tettay, Martynas Simanavičius, Milda Plečkaitytė, Lucas Hultgren, Anders P. Håkansson, Lars Malmström, Simon Ekström, Lotta Happonen, Johan Malmström

## Abstract

*Streptococcus pneumoniae* remains a major public health concern, largely due to the limited serotype coverage and other constraints of pneumococcal conjugate vaccines, as well as the increasing antimicrobial resistance among circulating strains. In the pursuit of protein-based pneumococcal vaccines, pneumolysin (PLY), a secreted multifunctional cholesterol-dependent cytolysin, represents a promising target. To date, the relationship between the B-cell epitope landscape and neutralising PLY-specific antibody responses has remained elusive, hindering the rational design of effective PLY-based vaccines. Using a panel of PLY-specific monoclonal antibodies, functional assays, multimodal protein mass spectrometry and data-driven computational modelling, we mapped the structural epitope landscape of native PLY and linked epitope-paratope interactions to neutralising potency. We further refined and structurally characterised a protective cross-species epitope conserved among homologous cholesterol-dependent cytolysins (CDC). This epitope provides a promising foundation for rational, epitope-focused vaccine design, offering a pathway toward species-independent vaccines targeting the CDC protein superfamily.

## Introduction

*Streptococcus pneumoniae* is a Gram-positive opportunistic bacterial pathogen that can cause infections such as pneumonia, septicaemia, and meningitis ^1^. The World Health Organisation has classified *S. pneumoniae* as a priority pathogen due to the high carriage rate in the human population, the underestimated disease burden and its role in causing over 800,000 deaths in children annually ^2^. Disease prevention can be achieved through the use of pneumococcal conjugate vaccines (PCVs), which utilise capsular polysaccharides as antigens ^3^. However, the overall effectiveness of PCVs is constrained by multiple factors, including limited serotype coverage of PCVs, the occurrence of serotype replacement, and the increasing antibiotic resistance among emerging strains ^4,5^. Moreover, polysaccharide antigens exhibit intrinsically weaker immunogenicity compared to protein-based immunogens ^6,7^.

Thus, protein-based pneumococcal vaccines, targeting conserved surface and secreted proteins are being explored as an alternative strategy ^8–10^. One promising target for both antibody therapy and vaccine design is pneumolysin (PLY), a secreted multifunctional toxin that plays an important role in pneumococcal pathogenesis ^11^. Combined with serotype independence, high level of carriage and conservation across pneumococcal strains, relatively safe immunisation profile together position PLY as a promising vaccine target ^12^. PLY is a canonical member of the cholesterol-dependent-cytolysin (CDC) protein superfamily found in numerous Gram-positive bacterial species, sharing the conserved four-domain structure, cholesterol binding, and oligomeric β-barrel pore formation ^13^.

The current state-of-the-art vaccine design for PLY relies on generating detoxified versions of PLY (dPLY) or subunit constructs, such as PlyD4, which focuses on domain 4 critical for cellular membrane recognition ^14,15^. However, the altered structure of dPLY/PlyD4 due to point mutation/truncation can inadvertently lead to non-functional antibody responses, combined with immunodominant sites embedded with full-length antigen during vaccination ^16^. Effective and protective vaccine-elicited immunity relies on antibodies targeting specific epitopes of biological relevance (*e.g.*, neutralisation), many of which are conformational and require preservation of the protein’s native three-dimensional structure ^17^. While PLY is highly conserved across *S. pneumoniae* strains, achieving broad cross-strain or even cross-species protection requires deciphering epitopes that elicit broadly neutralising responses. Together, these factors underscore the immediate need for high-precision epitope mapping, structure-guided design and antibody-guided vaccine design ^18,19^, thereby facilitating the rational design and development of the novel vaccine capable of inducing durable, high-magnitude, functionally relevant antibody responses ^20^.

A key element of epitope-focused vaccinology is the use of monoclonal antibodies (mAb) to identify protective antigen regions in their native three-dimensional conformations, which can then guide epitope selection and the structural redesign of new vaccine constructs. The growing availability of high-quality antigen-specific mAbs ^21^, now provides opportunities to extensively map structural epitope landscapes to better understand the correlates between epitope-paratope interactions and protective mechanisms. Achieving this requires methods for high throughput and high-fidelity epitope mapping, a persistent analytical challenge for bacterial antigens. Conventional techniques such as peptide array assays and alanine scanning mutagenesis, primarily identify linear epitopes and often fail to resolve conformational epitopes ^22^, which are critical for robust *in vivo* immuno-protection. Moreover, structural biology techniques (X-ray crystallography and NMR) are low throughput and demanding in time and cost (cryo-EM), which becomes increasingly problematic given the number of high-quality, antigen-specific mAbs to screen.

Mass spectrometry (MS) has emerged as a promising approach to define epitopes of relevance for immunity ^22–28^. Over the last decades, several different protein mass spectrometry techniques have proven useful for mapping epitopes in their native three-dimensional conformation including hydrogen/deuterium exchange MS (HDX-MS), chemical cross-linking MS (XL-MS), epitope excision/extraction MS and foot-printing labelling MS ^29–31^. Although these orthogonal methods generate informative data, they are also associated with certain limitations ^22^. Consequently, the complementary use of two or more of these methods, coupled with data-driven computational modelling is increasingly employed and has been shown to enable the successful characterisation of structural epitopes of functional relevance ^24,32–36^. When combined, protein MS methods can be applied to broader panels of mAbs, scaled up to more complex samples, and used to elucidate protein dynamics resulting from destabilisation and allosteric effects. These capabilities provide new possibilities to profile structural epitope landscapes for a given antigen targeted by multiple mAbs of various properties, thereby enabling a deeper understanding of the link between the formation of binding interface and neutralising potency.

In this study, we used a panel of anti-PLY mAbs, *de novo* MS sequencing, a combination of structural MS techniques, and computational analysis to profile the structural epitope landscape of PLY. The results demonstrated that distinct epitopes are linked to the functional properties of each mAb. Notably, we also determined a broadly protective B-cell epitope shared among CDCs from unrelated bacterial species, which possesses great potential for follow-up studies.

## Results

### No correlation was found between binding affinity and neutralisation level of the PLY-specific mAbs

High-resolution epitope mapping using a panel of mAbs can decipher and define epitopes of biological relevance for rational vaccine design. Here, we employed a multimodal MS workflow to map the epitope landscape of PLY using a panel of specific mAbs (**Fig. 1**). By integrating *de novo* MS sequencing, XL-MS, HDX-MS and data-driven computational modelling, we obtained detailed information for the antigen-antibody complexes, enabling the structural profiling of the epitope landscape of PLY under near-native conditions. Epitopes critical for neutralisation can serve as a starting point for the designing novel epitope-focused vaccines and guiding the development of direct anti-virulence therapeutic strategies (**Fig. 1**).

**Figure 1.**
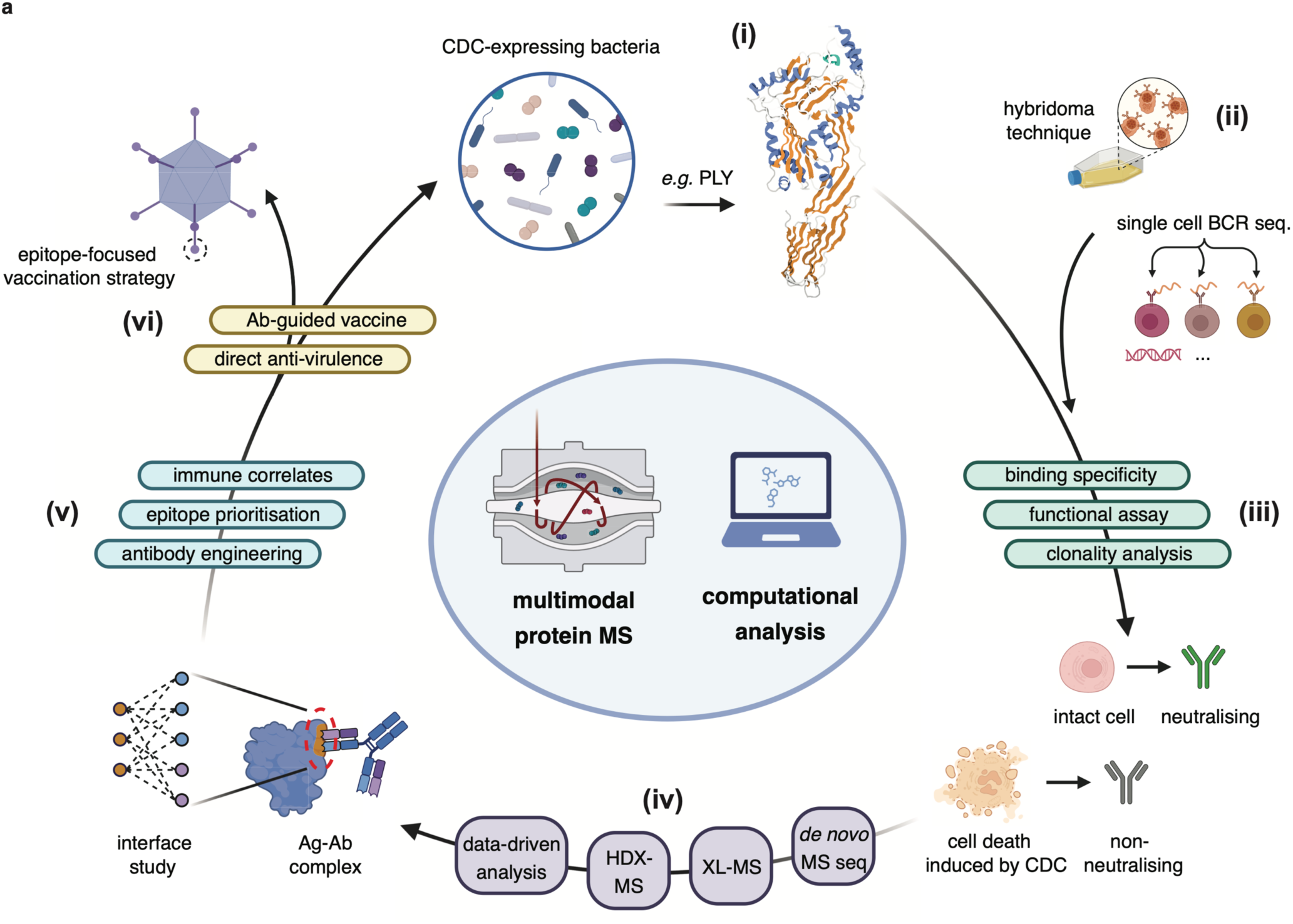
Multimodal mass spectrometry-powered integrative structural biology workflow to define the epitope landscape of a bacterial virulence factor. The illustration presents an overview of the multimodal MS (MMS) workflow used in this study. Many bacterial pathogens produce cholesterol-dependent cytolysins (CDC), a protein superfamily capable of lysing eukaryotic cells, with (i) pneumolysin (PLY) from *Streptococcus pneumoniae* serving as a representative CDC. Using PLY as a target, specific monoclonal antibodies (mAb) can be generated either by (ii) hybridoma technique or through single-cell B-cell receptor sequencing. The panel of mAbs are ranked based on (iii) binding affinities and their capacity to neutralise PLY-mediated cytolysis to prioritise the most effective clones. These antibody clones are subjected to (iv) *de novo* MS sequencing to obtain the primary structure, respectively. High-confidence structural models of the antigen-antibody complexes can be generated by structural mass spectrometry techniques, including (iv) XL-MS and HDX-MS, in combination with data-driven computational analysis, providing detailed information of the binding interfaces for each pair. These epitope-paratope interactions provide a mechanistic link between the (v) binding interfaces and neutralisation, serving as a starting point for (vi) rational design of novel epitope-focused vaccines and development of direct anti-virulence mini-binder therapeutics.

As a starting point, ten PLY-specific mAbs from the previous study were selected ^21^. These antibodies were generated using the hybridoma technique following immunisation of BALB/c mice with PLY. An indirect ELISA (**Fig. 2a**) was used to confirm the binding specificity of all mAbs to PLY. In parallel, we evaluated the ability of these mAbs to neutralise PLY-induced haemolysis, by incubating PLY with a serial dilution of each mAb. Diluted sheep erythrocytes were then added, incubated and pelleted down to harvest the supernatant, after which the absorbance of released hemoglobulin was measured to quantify neutralisation for each mAb (**Fig. 2b**). Among the ten mAbs, only seven are able to protect erythrocytes from PLY-induced haemolysis (**Fig. 2c, Extended Data Fig. 1a**). Of these, five clones, namely 3A9, 6E5, 3F3, 12F11, and 3C10, exerted significantly higher level of neutralisation, as indicated by the half-maximal inhibitory concentrations (IC_50_) from a two-fold serial titration (**Fig.2d, Extended Data Fig. 1a**).

**Figure 2.**
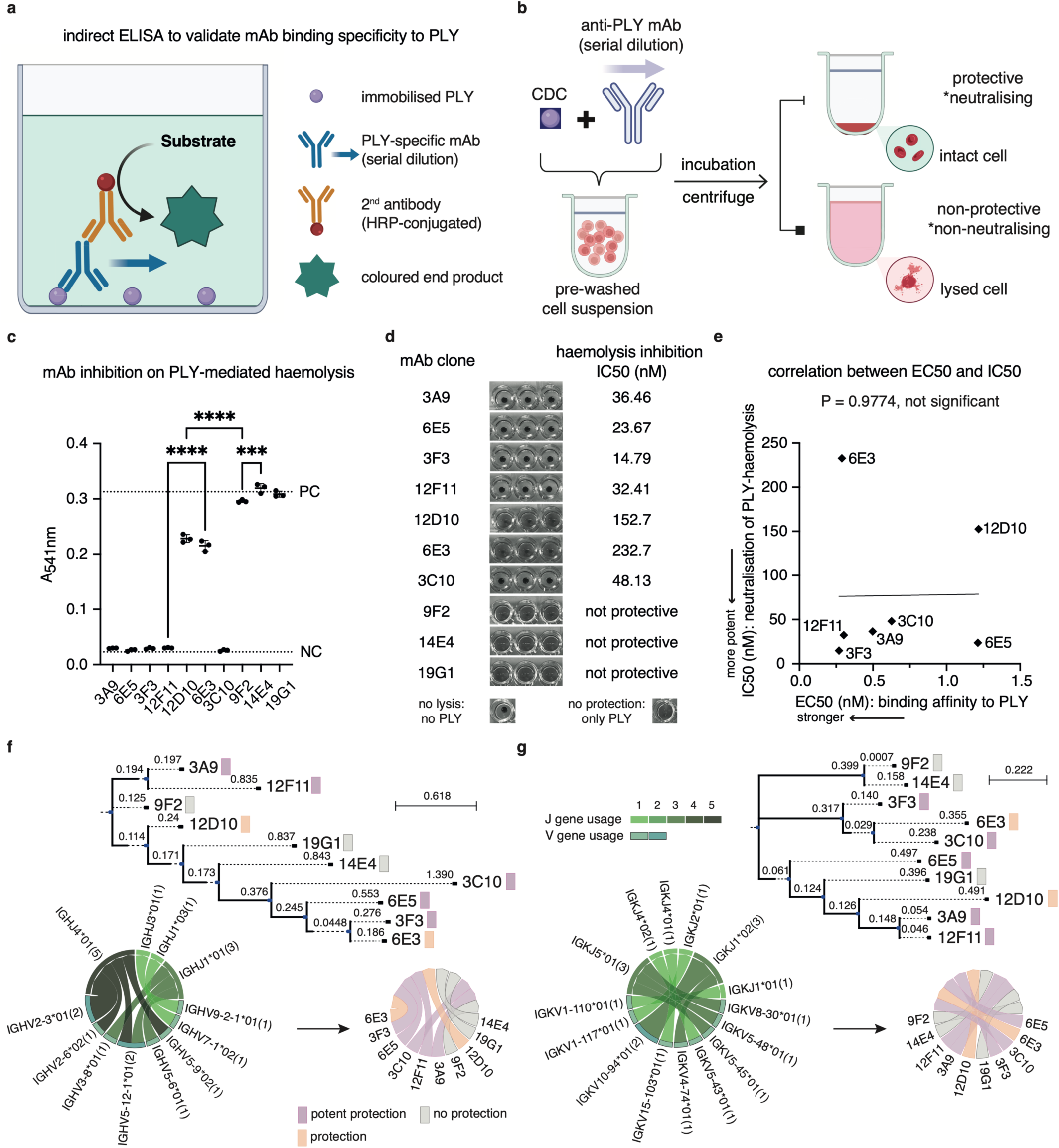
Characterisation of PLY-specific monoclonal antibodies and mAb clonal analysis. **a**) The ten mAbs were subjected to indirect ELISA to determine the binding avidity to antigen PLY. EC50 values, representing the half-maximal effective concentration for PLY-binding, were calculated from serially diluted mAbs. **b**) To determine the neutralisation level of the mAbs, PLY was added to diluted sheep erythrocyte (RBC) suspensions, pre-incubated separately with each mAb in in serial dilutions. After incubation and centrifugation, the absorbance of the supernatant was measured at 541 nm to quantify the amount of haemoglobin released due to cell lysis. **c**) Neutralisation capacity of PLY-induced haemolysis at the least diluted mAb concentration (533 nM) is shown. The positive control (PC): PLY without mAb addition. Negative control (NC): no PLY added. **d**) Monochromic images showing the level of haemolysis after pelleting the unlysed RBCs by centrifugation in triplicates. The table on the right displays the calculated half-maximal inhibition concentration (IC50) values. Representative PC and NC examples are shown at the bottom. **e**) Linear regression analysis of the obtained IC50 and EC values was performed for the seven mAbs with measurable IC50 values. All ten mAbs were subjected to *de novo* MS sequencing to construct B-cell lineage trees and analyse V-J gene pairwise preferences based on multiple sequence alignments (MSA) of **f**) heavy chain or **g**) light chain sequences. The tree topology shows distinct clades representing B-cell clones that have undergone somatic hypermutation. Branch lengths, as indicated by the scale, reflect the relative evolutionary distance, highlighting the accumulation of mutations during affinity maturation. V-J gene pairings are summarised in chord diagrams coloured by gene usage count, annotated with their respective germline gene names and level of protection (neutralisation).

To investigate whether the neutralisation potency was dependent on mAb binding affinity (avidity), a non-linear regression analysis was performed to determine the half-maximal effective concentration (EC_50_) value of each mAb (**Extended Data Fig. 1b**). All mAbs demonstrate strong binding to PLY, with EC_50_ values ranging from picomolar to nanomolar level. Linear regression analysis comparing the EC_50_ and IC_50_ values of the seven neutralising mAbs with measurable IC_50_ values (**Extended Data Fig. 1c**) revealed no correlation between apparent binding affinity and neutralisation capacity (**Fig. 2e**). Taken together, these results indicate that stronger antibody binding, though established by extensive affinity maturation *in vivo*, does not directly translate into effective or higher level of toxin neutralisation.

### *De novo* MS sequencing determined the primary structure of the PLY-mAbs

To determine the primary structures of the mAbs, multi-enzyme digestion coupled with *de novo* MS sequencing was applied. Five proteases, including trypsin, pepsin, chymotrypsin, elastase, and alpha-lytic protease, were used to generate overlapping peptides of varying length for each mAb, which were then subjected to *de novo* sequencing. The resulting sequenced peptides were assembled into full-length protein sequences for both the heavy and light chains, achieving 100% coverage for all included mAbs (five representative clones are displayed in **Extended Data Fig. 1d**-**h**). Structural models of the derived fragment antigen-binding (Fab) domains were subsequently predicted using IgFold ^37^ and AlphaFold-Multimer^38^ (0.648 ≥ pTM ≤ 0.88).

Reconstruction of B-cell lineage trees of the ten mAbs for both the heavy (**Fig. 2f**) and light chain sequences (**Fig. 2g**) revealed ten clonotypes with distinct maturation trajectories. Several mAbs shared pairings of V-J genes, particularly between the phyletic clones 3A9 and 12F11, also 3F3 and 6E3 (**Extended Data Table 1**). Identical V-J gene usage across both chains suggests that these mAbs may recognise the same antigenic region but differ in binding properties due to variations in their complementarity-determining region (CDR) loops. Branch lengths reflect the degree of divergence from the corresponding germline reference, with longer branches indicating a greater accumulation of mutations during somatic hypermutation (**Fig. 2f**-**g**). Overall, the heavy chains exhibited more extensive hypermutation than the light chains. These findings suggest that the epitope-binding specificities of these ten mAbs are shaped by unique pairing and combination of V-J gene pairings in combinations with distinct CDR loop architectures.

### Cross-linking mass spectrometry probed the epitope region targeted by different mAbs

To probe the antigen regions involved in neutralisation for these mAbs, we used cross-linking MS to detect unique intermolecular cross-links formed between residues positioned closely in three-dimensional space (**Fig. 3a**). Full-length PLY consists of three discontinuous domains (D1-3) and a C-terminal domain 4 (D4) (**Fig. 3b**). To map the proximal binding interfaces, each mAb was cross-linked to PLY using two different homobifunctional chemical linkers (DSS, disuccinimidyl suberate; DSG, disuccinimidyl glutarate) with different spacer arm length (11.4 Å; 7.7Å). Interprotein cross-linked peptide pairs were identified and searched using tandem MS.

**Figure 3.**
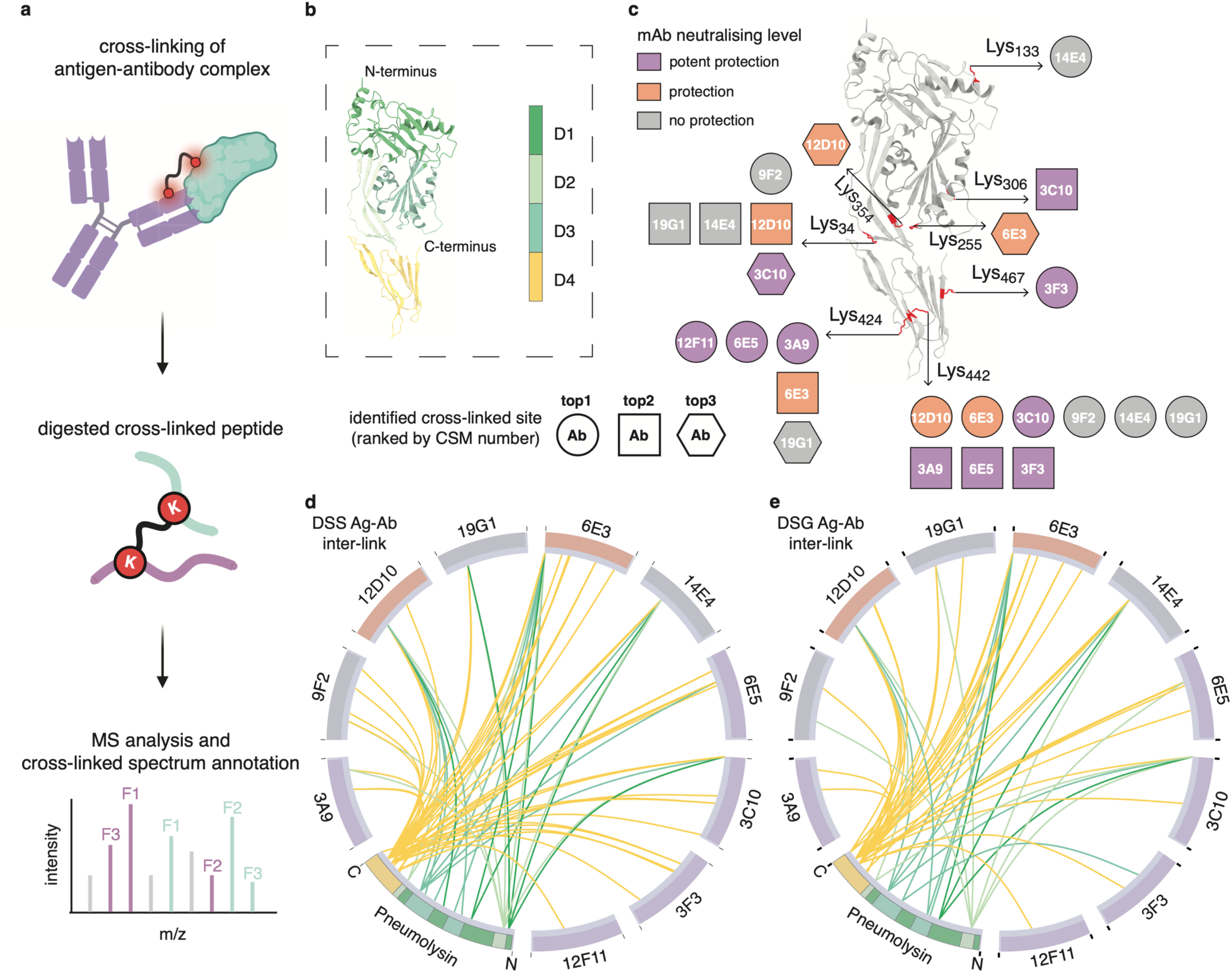
Cross-linking mass spectrometry analysis of different pneumolysin-mAb complexes. To determine reactive residues located within or in proximity to the binding interfaces in the antigen-antibody complexes, cross-linking MS (XL-MS) was performed. **a**) Either DSS- or DSG-duplet cross-linkers were used to cross-link PLY with each of the ten intact mAbs, separately. Mass spectrometry analysis and identification of various interprotein cross-linked peptides confirmed the formation of antigen-antibody immune complexes. **b**) The structural annotation of full-length PLY is shown, coloured according to domains. **c**) The three most frequently detected cross-linked peptides between different antibody clones and PLY were summarised, ranked by the count of cross-linked spectrum matches (CSM) and mapped onto the antigen structure. Cross-linked lysine (Lys) residues are labelled with the residue number, highlighted in red and displayed in side-chain stick representation. Antibody clones are coloured according to their measured neutralising potency. **d**-**e**) Two circular plots summarise all unique hetero-interprotein cross-links identified between PLY and the corresponding mAb (heavy chain and light chain overlaid), colour-coded by neutralisation potency. Cross-link edges are coloured according to the PLY domain involved.

The identified cross-linked sites on the PLY were sorted by the number of cross-linked spectrum matches (CSM) for each PLY-mAb pair. For the neutralising clones 3A9, 6E5, and 12F11, the top cross-linked site was Lys_424_ in D4, and Lys_467_ in D4 for the neutralising clone 3F3 (**Fig. 3c**). In contrast, the Lys_442_ in D4 was the most frequently detected cross-link for clones 12D10, 6E3, 3C10, 9F2, 14E4, and 19G1. For several of the latter clone groups, top 2 or top 3 cross-linked sites were found far from the D4 in space including 12D10, 14E4 and 19G1 (Lys_34_ in D2), 6E3 (Lys_255_ in D3) and 3C10 (Lys_306_ in D3). These clones had no (9F2, 14E4, 19G1) or lower (12D10 and 6E3) neutralisation effect, demonstrating that epitopes located to D4 is primarily linked to neutralisation. Overall, we identified 53 unique DSG cross-links from 296 CSMs (**Fig. 3d**), and 65 unique DSS cross-links from 192 CSMs (**Fig. 3e**), within 20 ppm MS/MS fragments mass accuracy (**Extended Data Fig. 2a-j**). In conclusion, XL-MS was able to confidently identify cross-linking sites between PLY and all ten mAbs, linking highly neutralising mAbs to epitope regions predominantly located in D4 of PLY.

### Hydrogen/deuterium exchange mass spectrometry determined multiple epitopes on PLY

To investigate the epitopes for five selected mAbs, we performed HDX-MS. In this approach, surface residue hydrogens are exchanged with deuterium, and subsequent MS analysis detects mass shifts between different states, revealing protein binding interfaces and providing unique insight into in-solution protein dynamics (**Fig. 4a**).

**Figure 4.**
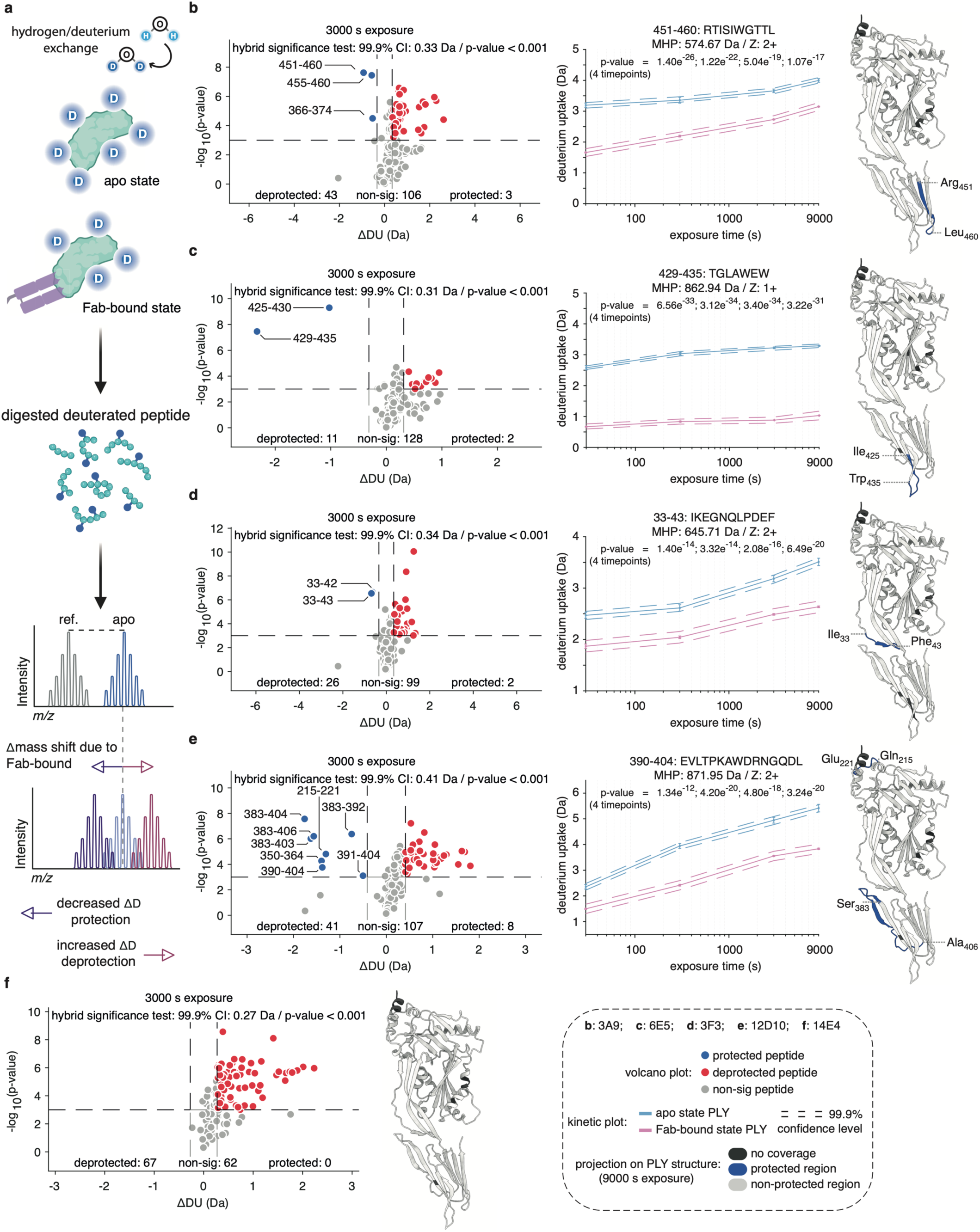
Hydrogen/deuterium exchange mass spectrometry analysis of pneumolysin protein during five different Fab bindings. To further refine the epitope regions determined by XL-MS, HDX-MS experiments were performed on PLY in both the apo state (PLY alone) and in the complex state (Fab-bound) **a**) In the HDX-MS experiment, mass shifts of pepsin-digested peptides, resulting from changes in deuterium uptake among different states, are measured by the mass spectrometer to define interaction interfaces and assess protein in-solution dynamics. **c-f)** In the complex state, excessive Fab derived from the mAbs **b**) 3A9, **c**) 6E5, **d**) 3F3, **e**) 12D10, and **f**) 14E4, was used to determine binding interfaces and to assess protein dynamics of PLY in PBS solution upon Fab binding. A global-level hybrid significance test was applied to determine protected, non-significant, and deprotected peptides across four different deuteration intervals (30, 300, 3000, and 9000 s). Kinetic plots of representative protected peptides are shown, with significance levels determined by multiple linear regression analysis. Significantly protected peptides identified at the longest deuteration time (9000 s) were mapped onto the PLY structure and highlighted in blue.

Fab fragments were generated from the five selected mAbs, three strongly neutralising (3A9, 6E5 and 3F3), one weakly neutralising (12D10), and one non-neutralising clone (14E4). Excessive Fab was used to saturate the PLY binding and to form PLY-Fab complexes. Both the apo state and the Fab-bound state samples were subjected to deuterium labelling over time from 30 to 9,000 seconds, to define the structural epitopes and to assess PLY protein dynamics upon Fab binding. The hydrophobicity (lipophilicity) of each residue was calculated (**Extended Data Fig. 3a**) as hydrophobic regions typically exchange less frequently with solvent during deuterium labelling, limiting the ability to assess protein dynamics in certain regions. The overall sequence coverage ranged from 96.6% to 97.7%, essentially encompassing all structured regions (**Extended Data Fig. 3b**).

The HDX-MS analysis enabled confident identification of the PLY epitopes for four out of the five Fabs. For these four clones, proximal inter-links were also identified, corroborating the epitope peptides defined by HDX-MS. Three clones recognise epitopes in domain 4, where 3A9 primarily binds to the D4 Loop 1 region Arg_451_-Leu_460_ (**Fig. 4b**), 6E5 to the D4 undecapeptide Ile_425_-Trp_435_ (**Fig. 4c**), and 12D10 to the D4 Ser_383_-Ala_406_ (**Fig. 4e**). As shown in **Fig. 2** previously, 3A9 and 6E5 are strongly neutralising clones whereas the 12D10 exhibited a lower degree of neutralisation. On the other hand, the neutralising 3F3 recognised an epitope in D2 Ile_33_-Phe_43_ (**Fig. 4d**). No epitope information could be confidently determined for the 14E4 (**Fig. 4f**).

Interestingly, there was a pronounced deprotection pattern in part of D3b upon Fab binding (**Extended Data Fig. 3c**). This phenomenon may result from binding-induced destabilisation or allosteric effects. These results demonstrate that epitopes with different locations can lead to comparable levels of neutralisation and that not all epitopes in D4 confer optimal protection. Further analysis is required to clarify how these distinct deprotection profiles on Fab-bound PLY correlate with biological outcomes. In summary, combining multimodal protein MS, we identify parts of structural epitope targeted by four distinct mAbs, and successfully connect epitope specificities to the neutralising potency of mAb against PLY haemolysis.

### Structural mass spectrometry on 6E5-CDC immune complexes identified a cross-species broadly neutralising epitope

PLY is part of the CDC superfamily of β-barrel pore-forming exotoxins, which share similarities in sequence, 3-D structure, and biological function. Previously, clone 6E5 has been shown to cross-react with other CDCs ^21^, but no specific epitope could be defined. To further investigate the cross-reactivity mechanism, an indirect ELISA was performed to confirm that 6E5 binds equally strongly to all included CDCs except for SLO, where the interaction was weaker (**Fig.5a**). Haemolysis inhibition assays with PLY, PFO, and SLO confirmed that 6E5 effectively neutralised erythrocyte lysis induced by all three toxins, including SLO (**Extended Data Fig. 4a**-**b**). These findings suggest that the neutralising epitope targeted by 6E5 is expected to be both physiochemically and conformationally conserved across the CDC protein superfamily.

To determine the location of the epitope recognised by 6E5 in other CDCs, XL-MS experiments were repeated using two linkers to cross-link the 6E5 Fab to each of the six CDCs (PLY, PFO, ILY, VLY, INY, and LLO). Inter-links supported by multiple CSMs were found in all CDC-Fab complexes (**Fig. 5b**). For PLY, the dominant cross-linked site was Lys_424_, consistent with the previous PLY-mAb 6E5 XL-MS experiments, located near the tip of D4 (**Fig. 5c**). This site aligns with the epitope determined by HDX-MS (PLY peptide, Ile_425_-Trp_435_). For the other CDCs, the most frequently detected cross-linked sites were: Lys_482_ of ILY, Lys_473_ of VLY, Lys_472_ of INY, Lys_435_ of PFO, and Lys_460_ of LLO (**Fig. 5c**). All these residues occupy structurally analogous position to PLY Lys_424_ within their respective CDC structure, supporting that the epitope recognised by 6E5 is structurally conserved in D4 across generic CDC toxins. To validate these findings, HDX-MS was performed on the PFO, produced by *Clostridium perfringens*, to determine the PFO epitope targeted by 6E5, achieving 100% sequence coverage of PFO between states (**Extended Data Fig. 4c**). HDX-MS successfully identified one of the peptides (**Fig. 5d)** with a significant change in deuterium incorporation, consistently demonstrating protection (**Extended Data Fig. 4d**), across the labelling intervals (**Fig. 5e**). These results experimentally confirms that the cross-reactive epitope is located at the tip of D4 and overlaps with undecapeptide of PFO (**Fig. 5f**). Notably, a global deprotection pattern was also revealed in D3b region of PFO during 6E5 Fab binding (**Extended Data Fig. 4d**), analogous to the observations made for PLY-6E5 interaction.

**Figure 5.**
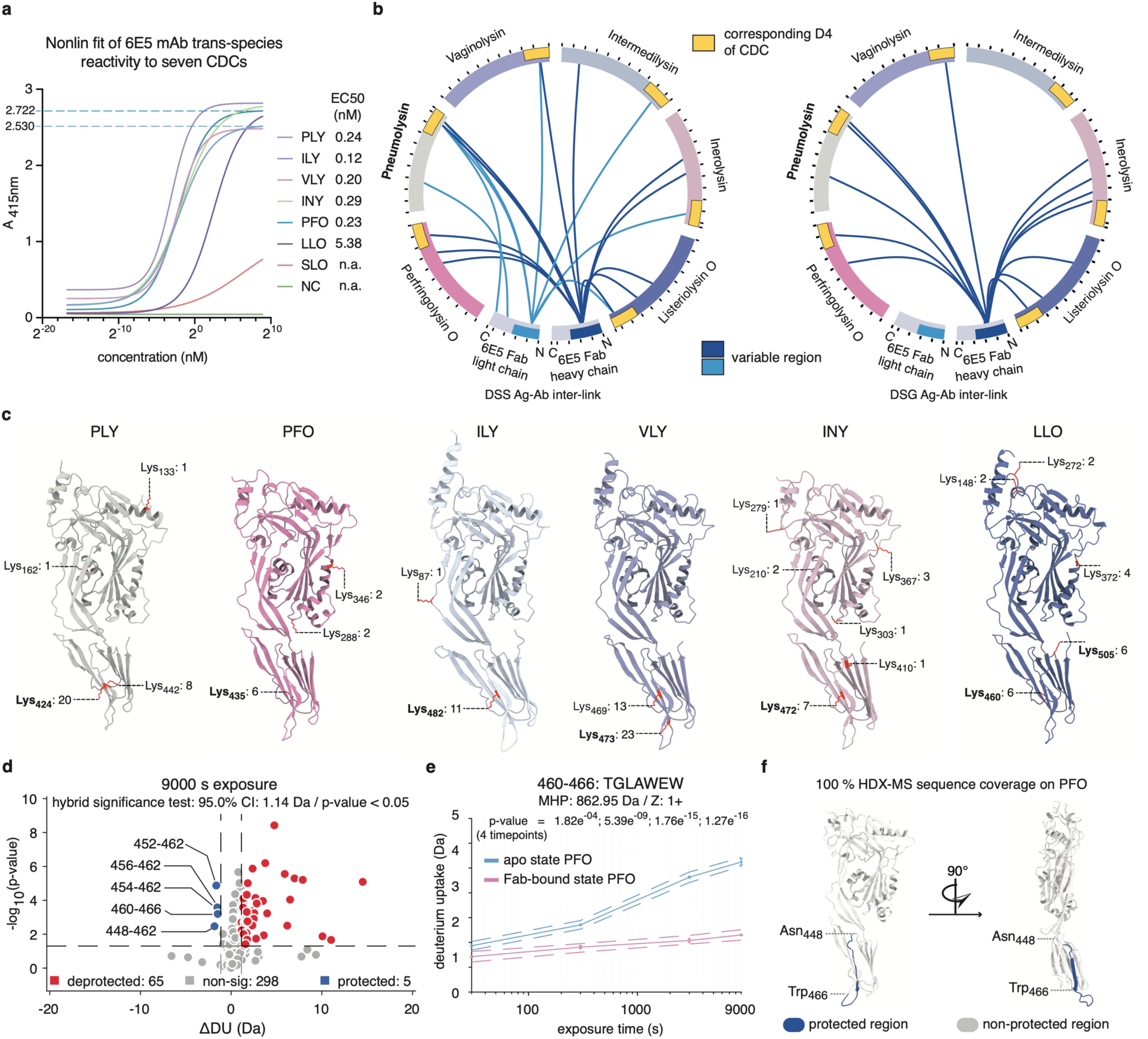
Combined structural mass spectrometry analysis of the cross-species reactive antibody 6E5. **a**) Indirect ELISA was repeated to measure the binding specificity of antibody clone 6E5 to several unrelated CDCs from other bacterial species, including intermedilysin (ILY), vaginolysin (VLY), inerolysin (INY), perfringolysin O (PFO), listeriolysin O (LLO), and streptolysin O (SLO). EC50 values were calculated by non-linear regression analysis for each CDC-6E5 pair, with the maximum binding site indicated by a dashed line for PLY and PFO. **b**) 6E5 Fab domain was cross-linked to each of the six CDCs using either DSS- or DSG-duplet cross-linkers. The circular plots summarise unique inter-protein DSS- and DSG-linked sites identified between 6E5 Fab and the corresponding CDCs, colour-coded by CDC. Domain 4 (yellow boxes), and the variable regions (blue boxes) of the heavy/light chains are highlighted. Cross-link edges are coloured by different antibody chains. **c**) All cross-linked sites from the DSS- and DSG-XL datasets combined, are shown on the CDC models as sidechain stick representations, annotated with the reported CSM counts and the most frequent site(s) highlighted in bold. Next, HDX experiment was performed on the apo PFO and the PFO in complex with 6E5 Fab. A global-level hybrid significance test was applied to determine PFO peptides with differential deuterium uptake across different deuteration intervals between states. **d**) The volcano plot displays the PFO peptides with a significant change in deuterium uptake, due to 6E5 Fab interaction, after a 9000 s deuterium exposure. **e**) A kinetic plot of one representative protected peptide with significance levels determined by multiple linear regression analysis. **f**) The protected regions identified at the longest deuteration exposure (9000 s) were mapped onto the PFO structure from two viewpoints and highlighted in blue.

### High-quality data-driven modelling revealed 6E5-bound interfaces on PLY and PFO

The antigen-antibody (Ag-Ab) binding interface typically covers a total surface area of approximately 800-2,000 Å^2^ ^39–41^. Accurate modelling of Ag-Ab pairwise complex, however, remains challenging when few or no experimental distance constraint data are available ^42^. To overcome this, we implemented HADDOCK (High Ambiguity Driven protein-protein DOCKing), incorporating the multimodal experimental restraints from XL-MS inference and peptide-level HDX-MS distance constraints for the antigen, and refining the Fab search space by the CDR HV loops only found protected in HDX-MS. Prevalent XL-MS cross-links were applied as centre-of-mass restraints to constrain the search (**Fig. 6a**). The performance of the docking pipeline was benchmarked using an external dataset, which was able to generate a near-atomic fidelity modelling of the Ag-Ab (Fv) complex (**Supplementary Data Fig. 1a**), featuring both its global conformation and residue-level recall of the epitope (96%-100%), which was in line with the reference co-crystal structure ^43^.

**Figure 6.**
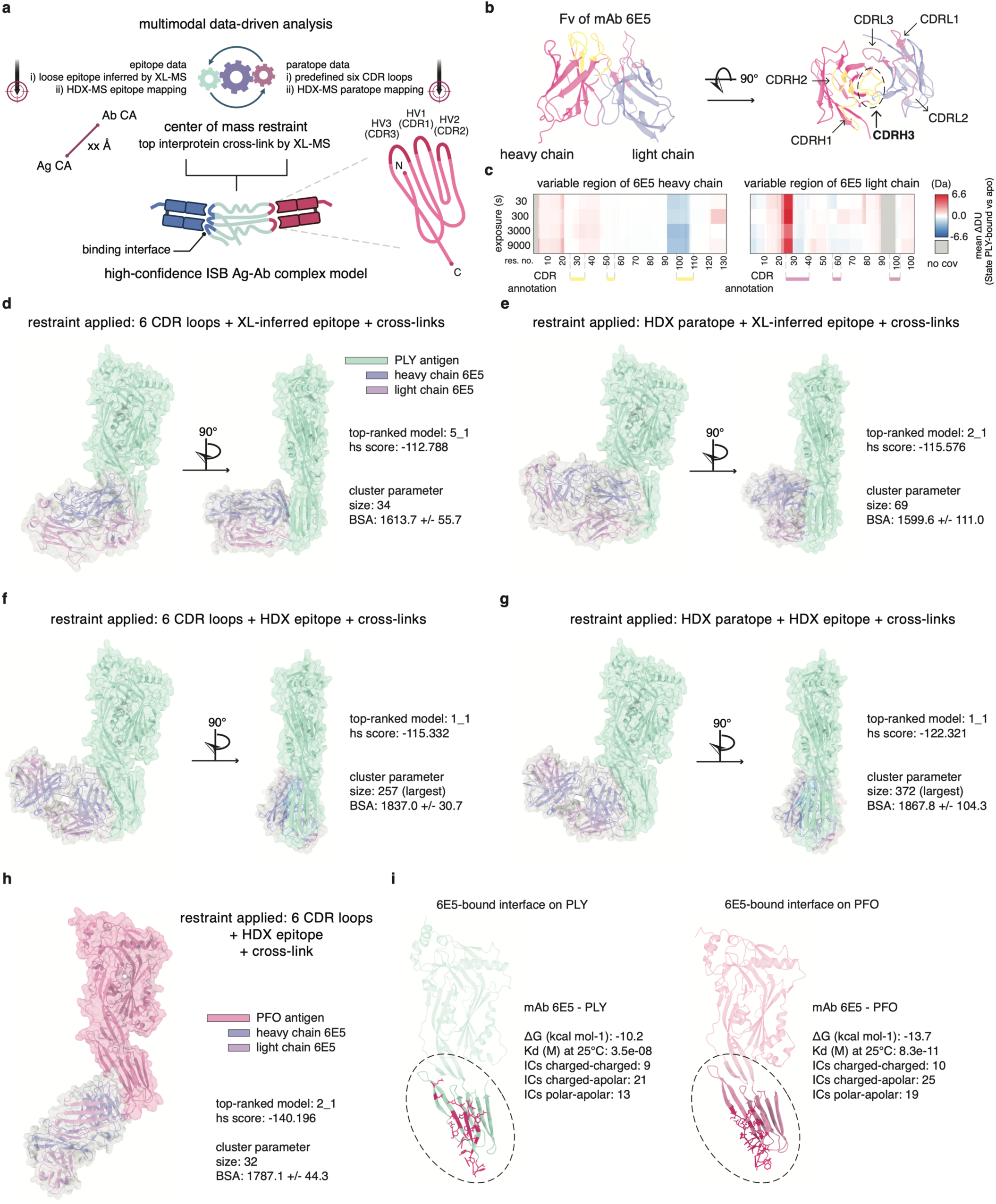
Data-driven computational modelling of various 6E5-CDC immune complexes. **a**) HADDOCK data-driven modelling of 6E5-CDC pair complexes was guided by multimodal restraints derived from XL-MS, HDX-MS, and the six pre-defined complementarity-determining regions (CDR) loops. **b**) An Fv (variable region of the Fab domain) model of 6E5 with the six CDRs labelled and colour-coded. **c**) HDX-MS was conducted on apo 6E5 and PLY-bound 6E5 identified paratope regions. The mean change of deuterium uptake per residue was plotted for variable regions of the 6E5 Fab, with aligned CDRs labelled across the protein sequences. **d**-**g**) Four docking scenarios were constructed based on epitope data (cross-link inferred vs HDX-determined) and CDR data (pre-defined vs HDX-determined). Prominent cross-links were imposed as centre-of-mass restraints. The docking process involved an initial 10,000 rigid-body docking, followed by a selection of 500 models for semi-flexible and final refinement. Four top models from each scenario, ranked by combined score (hs score), are shown from two viewpoints, annotated with cluster-wise size and buried surface area (BSA). **h**) An immune complex model of 6E5-PFO was constructed using the restraints and modelling procedure equivalent to **Figure 6f**. The top-ranked model is shown. **i**) The cross-species epitopes recognised by 6E5 on PLY and PFO were extracted from **Figure 6g** and **Figure 6h**, respectively. Interface residues were computed using PRODIGY and highlighted side-by-side with side chain stick representation in hot pink.

To define the PLY-interacting loops out of the six pre-defined CDRs of 6E5 (**Fig. 6b**), we first performed HDX-MS on the 6E5 Fab domain with or without the cognate antigen PLY, resulting in 88% sequence coverage of the Fab region (**Extended Data Fig. 5a**). CDRH3 exhibited significant protection consistently across all labelling intervals (**Fig. 6c**), identifying it as the principal contact region (**Extended Data Fig. 5b**), whereas CDRL1 displayed a more modest but still reproducible level of protection (**Extended Data Fig. 5c**).

Based on the available XL-MS and HDX-MS data, we modelled the PLY-n6E5 complex using four distinct restraint regimes: i) XL-MS inferred epitope (**Extended Data Fig. 5d, Fig. 6d**); ii) the same inferred epitope plus HDX-MS-confined CDR loops (paratope) (**Fig. 6e**); iii) HDX-MS-mapped epitope peptides (**Fig. 6f**); and iv) HDX-MS restraints on both epitope and paratope (**Fig. 6g**). The two most frequently observed intermolecular cross-links were used as centre-of-mass constraints in every run.

As the restraint density increased from (i) to (iv), the cluster containing the top-ranked model captured a larger fraction, the refined interface became more coherent, and both the overall hs score (HADDOCK scoring matric combining van der Waals, electrostatics, desolvation, and restraint energies) and buried surface area (BSA) matrices improved. Across the four scenarios, the computed cluster-wise BSA increased from ∼1600 Å^2^ to ∼1900 Å^2^.

We next modelled the 6E5-PFO (from *Clostridium perfringens*) complex using scenario (iii) including HDX-MS-mapped epitope peptides plus the dominant intermolecular cross-link (**Fig. 6h**). The top-ranked model positions 6E5 squarely over PFO domain 4, fully sequestering the undecapeptide motif. This model indicates that 6E5 inhibits PFO-induced haemolysis via the same mechanism as the neutralisation of PLY (**Fig. 6i**). Interface analysis revealed 54 inter-molecular contacts (ICs) between PFO and 6E5, exceeding the 43 ICs observed in the PLY–6E5 complex (**Fig. 6i**). This denser contact network indicates a more stabilised binding interface on PFO which aligns with the higher maximum binding site (Bmax) measured for PFO relative to PLY in the indirect ELISA assays (**Fig. 5a**).

### Summary of the epitope-function correlate of PLY domain 4 and the cross-species binding mechanism of 6E5

A common feature of all CDC proteins is the three loops and an extended undecapeptide located at the tip of D4 (as illustrated in **Fig. 7a**). The neutralising mAb 3A9, 6E5 and 12D10 all bound to PLY domain 4 with high affinity but targeted distinct epitopes. Clones 3A9 and 6E5 mAbs demonstrated higher neutralising potency compared to 12D10 (**Fig. 7b**). Both 3A9 and 6E5 recognise epitope peptide at the very tip of D4 with 3A9 targeting the L1 loop and 6E5 binding to the undecapeptide. Targeting these epitopes explains why these mAbs inhibit particularly PLY-cholesterol interactions as shown previously (**Fig. 7b**) ^21^, thereby preventing PLY attachment to the cellular membrane during the initial phase of cytolysis. In the same study, 6E5 was also found to specifically block PLY binding to the mannose receptor C-Type 1 (MRC-1) ^21^, an important interaction linked to pneumococcal intracellular survival ^44^. Our identification of the undecapeptide as the epitope for 6E5 corroborates previous findings that this motif is essential for MRC-1 recognition ^45^. In contrast, the 12D10 bound to an epitope higher up in the D4 and was associated with weaker neutralisation without inhibition of either cholesterol or MRC-1 binding (**Fig. 7b**) ^21^.

**Figure 7.**
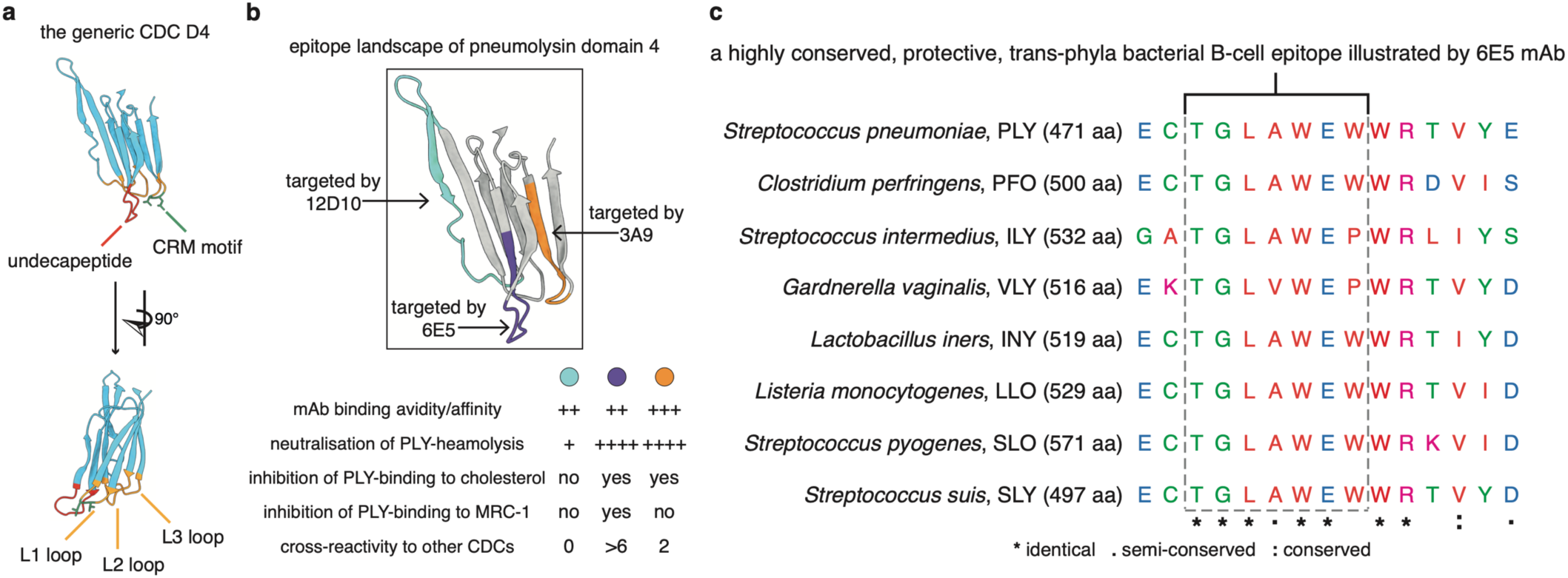
Characterisation of the broadly protective epitope targeted by cross-species neutralising 6E5. **a)** Loops L1 to L3, including the cholesterol-recognition motif (CRM), together with the undecapeptide, are highlighted in a generic domain 4 of CDC. Residues comprising the CRM and the undecapeptide are coloured in green and red respectively. **b**) Illustration of three core epitope peptides in PLY domain 4 identified in this study, colour-coded by the corresponding mAb clone. The functional implication of these binding sites is summarised below based on results derived from current and previous studies. **c**) CDC D4 sequence alignment highlights the region targeted by the cross-species reactive mAb 6E5, marked in dashed box (-TGLAWEW-), located within the highly conserved undecapeptide (-ECTGLAWEWWR-). Identical residues are marked with “*”, semi-conserved with “.” and conserved with “:”.

Although there is substantial sequence variation in D4 among different CDCs, the core epitope peptide recognised by 6E5 (Lys_429_-Lys_435_) is conserved across all known CDCs (**Fig. 7c**). Multiple sequence alignment, combined with the consistent identification of experimental inter-links between 6E5 Fab and various CDCs, suggests that antibodies targeting this structural epitope will neutralise all secreted CDCs by blocking the membrane binding of CDC. This epitope information provides an invaluable and rational starting point for design of broad CDC-specific inhibitors and for development of an epitope-focused vaccination strategy capable of neutralising CDCs across multiple species.

## Discussion

In this study, we combined multimodal protein MS with computational modelling to systematically characterise the structural epitope landscape of PLY. All tested mAbs exhibited strong affinity or avidity, reflecting extensive somatic hypermutation during *in vivo* affinity maturation. Of the ten mAbs, only seven neutralised PLY, with no clear correlation found between binding strength and neutralisation potency. Epitope mapping revealed that neutralisation was associated with epitopes located in D4, although not all epitopes within D4 conferred optimal protection, and epitopes in domains other than D4 could yield comparable levels of neutralisation. Two potently neutralising mAbs, 3A9 and 6E5, targeted epitopes at the very tip of D4, including the L1 loop region and the undecapeptide. The mAb 6E5, targeting the undecapeptide, blocked PLY binding to both cholesterol and MRC-1 and was linked to broad neutralisation across homologous CDCs produced by other bacterial species.

These findings underscore that the current pneumococcal vaccine design using full-length dPLY toxoids for immunisation is associated with two intrinsic shortcomings. First, full-length PLY harbours immunodominant epitopes that give rise to antibody clones with low or no neutralisation potency. Second, detoxification of PLY frequently involves the introduction of mutations into the undecapeptide motif to abolish cytotoxicity, thereby altering the native epitope integrity known for protective immunity across basically all CDC proteins ^46,47^. Together, our findings demonstrate that precise structural epitope mapping can successfully link epitope-paratope interactions to neutralising potency, providing a starting point for structure-informed, data-driven, and epitope-focused vaccine design strategy for complex bacterial pathogen ^48^.

Compared to previous attempts to map the epitope landscape on PLY, the multimodal approach used here enabled the successful identification of conformational epitopes. Earlier work identified only a single linear epitope peptide, located in domain 1 of PLY, recognised by 14E4 ^21^, with other epitope mapping limited by challenges inherent to overlapping PLY fragments and linear peptide screening approaches. By determining epitopes under near-physiological conditions here, with both antigen and antibody in their native three-dimensional conformation, structural epitopes can be defined more accurately. Unlike conventional approaches dependent on various derived fragments or peptides, structural MS-based methods preserve native antigen-antibody interfaces, with the correct conformation, while also providing unique structural insights and complementary information (*e.g.* transient interaction, protein in-solution dynamics) in addition to advanced biophysics techniques like crystallography and cryo-EM ^49–51^. Moreover, the integration of orthogonal MS techniques allows mutual corroboration and compensates for the limitations of each individual method ^22^. For example, HDX-MS is limited by low exchange rates in hydrophobic interfaces, as observed with clone 14E4, whereas XL-MS may fail to capture cross-links due to conflicting restraints or a lack of reactive residues near the formed epitope-paratope interface. Here, the data-driven modelling also benefits from integrating multimodal experimental data, as evidenced by consistently improved scoring metrics with increasing density of the restraints applied.

The protective epitopes identified here could serve as the basis for “hotspots” to guide the *de novo* design of broadly neutralising peptide/mini-protein binders and nanobodies. The XL-MS, HDX-MS and sequence alignment consistently pinpointed the cross-reactive protective epitope targeted by 6E5 to the undecapeptide region of PLY D4, which is highly conserved across nearly all reported CDC protein superfamily. A recent study reported a trans-phyla 15-mer immunodominant T-cell epitope spanning residues 427-441, although the correlation with *in vivo* protection has not yet been fully understood ^52^. Notably, this T-cell epitope overlaps in primary structure with the core B-cell epitope 429-435 targeted by 6E5. Emerging vaccine modalities, such as the two-component protein nanoparticle ^53–55^, could integrate cross-reactive and functional T-cell epitopes with conformational B-cell epitopes, providing a promising strategy for a serotype-independent vaccine to prevent pneumococcal colonisation and invasive infection.

## Methods

### Recombinant protein production and purification

All CDCs used in this study were recombinantly expressed and purified in accordance with previously reported protocols ^56^. The ten monoclonal antibodies (mAbs) were generated via hybridoma technology and purified as described ^21^. For Fab generation, five mAbs (3A9, 6E5, 3F3, 12D10, and 14E4) were processed using either a Mouse IgG1 or a generic Mouse IgG Fab Generation and Purification Kit (Thermo Fisher Scientific). The resulting Fab samples were buffer-exchanged into PBS using 30 kDa molecular weight cut-off (MWCO) centrifugal filters (Amicon, Merck) and concentrated to 4-5 mg/mL, as measured by a NanoDrop spectrophotometer (DeNovix). The sequence integrity of each Fab was subsequently verified by bottom-up mass spectrometry.

### *De novo* MS sequencing and BCR repertoire analysis

The ten mAbs were initially reduced in a solution containing 8 M urea, 100 mM ammonium bicarbonate, and 5 mM TCEP (Sigma Aldrich). Reduction was performed for 1 hour at 37 °C under continuous agitation (500 rpm) in a Thermomixer (Eppendorf). Alkylation was then conducted by adding 10 mM iodoacetamide (Sigma Aldrich) and incubating for 30 minutes in the dark at room temperature. Next, additional 100 mM ammonium bicarbonate was added to dilute the urea concentration to below 1 M. Four proteases (*i.e.*, trypsin, chymotrypsin, elastase, and alpha-lytic protease) were freshly reconstituted in their respective activation buffers, then added to each mAb sample at an enzyme-to-substrate ratio of 1:10 (w/w). The digestion proceeded overnight (16-18 hr) at 37 °C and 500 rpm, after which it was quenched by the addition of 40 µL of 10% formic acid. For pepsin digestion, the mAb samples were directly mixed with dissolved pepsin at a 5:1 (w/w) ratio in pH 2 buffer containing 0.01% TFA and 1 mM HCl. Following a 45-minute incubation at 37 °C and 500 rpm, the digestion was stopped by heating the samples to 95 °C. All digested samples were then purified using C18 reversed-phase micro spin columns (Harvard Apparatus). The cleaned-up peptide mixtures were concentrated in a SpeedVac (Eppendorf) to full dryness, and resuspended in buffer A (2% acetonitrile, 0.2% formic acid) prior to mass spectrometric analysis.

Approximately 1 µg of each protease-digested mAb sample, quantified by NanoDrop, was injected in triplicate onto an Ultimate 3000 UPLC system (Thermo Fisher Scientific) coupled to a Q Exactive HF-X Hybrid Quadrupole-Orbitrap mass spectrometer (Thermo Fisher Scientific). The peptides were concentrated and subsequently separated on a 50 cm µPAC Neo HPLC column (Thermo Fisher Scientific), following the manufacturer’s instructions. Two solvent mobile phases were used: solvent A (0.1% formic acid) and solvent B (0.1% formic acid, 80% acetonitrile). A linear gradient from 4% to 38% B was applied over 120 minutes at a flow rate of 450 nL/min.

Data were acquired in data-dependent acquisition (DDA) mode, starting with an MS1 scan from m/z 300 to 2,000 at a resolution of 120,000, an automatic gain control (AGC) target of 3e6, and a maximum injection time of 100 ms. The top 20 precursors were selected for MS2 scans at a resolution of 30,000, an AGC target of 1e5, a 20 ms injection time, and stepped normalised collision energies (NCE) of 20, 26, and 35. Charge states of 1, 6-8, and above were excluded, except for samples digested with enzymes other than trypsin or pepsin, where singly charged ions (1+) were included. LC-MS/MS performance was regularly evaluated by injecting and analysing a yeast protein extract digest (Promega) for quality assurance. In total, 150 DDA datasets were acquired and analysed using the Supernovo *de novo* MS sequencing algorithm (Protein Metrics) with the monoclonal antibody sequencing and assembly workflow.

Following confirmation of each clone’s heavy-and light-chain sequences, Abalign ^57^ was employed for sequence alignment and repertoire analysis. Framework and CDRs were defined accordingly to the Chothia canonical numbering scheme. B-cell lineage trees were then reconstructed from the variable domains of either the heavy or light chains using FastTree2 method ^58^, with the branch lengths normalised to represent the degree of affinity maturation tracks.

### Indirect ELISA and haemolysis inhibition assay

For the indirect ELISA, equivalent amounts of tested CDC antigens (PLY, ILY, VLY, INY, PFO, LLO, and SLO) were coated onto MaxiSorp plates (Thermo Fisher Scientific) and incubated at 4 °C overnight. Plates were washed three times with PBST (PBS containing 0.05% Tween 20) between each step. Unspecific binding sites were blocked by incubating with freshly made 2% (w/v) bovine serum albumin (BSA) solution. Each monoclonal antibody (mAb) clone was then added to the plates in a five-fold dilution series, starting with an antigen-antibody molar ratio of 2:1. After incubation, a goat HRP-conjugated anti-mouse IgG secondary antibody was applied at a 1:3000 dilution and incubated for 1 hour. Then, a freshly prepared HRP substrate solution was introduced, and after 5 minutes of colour development, the reaction was stopped with 50 mM H₂SO₄. Absorbance was measured at 415 nm wavelength, and antibody binding affinity/avidity to the specified antigen was inferred as the EC_50_, representing the half-maximal effective concentration (calculated from the dilution series) required to achieve maximal reactivity.

For the CDC-haemolysis inhibition assay, 80 ng of each CDC toxin (PLY, PFO, and SLO) was incubated with each mAb clone in a two-fold dilution series, starting from an antibody concentration equivalent to 4 µg of protein. The CDC-mAb mixtures were incubated for 30 minutes at 37 °C, after which freshly washed (3x) and diluted (4% v/v) sheep erythrocyte suspension (Fisher Scientific) was added. The samples were then incubated for an additional 60 minutes at 37 °C. Following incubation, the plates were centrifuged, and the supernatants were carefully harvested and transferred to new plates for absorbance measurement at 541 nm wavelength. Antibody neutralising potency against CDC-induced haemolysis was determined as the IC_50_, representing the half-maximal inhibitory concentration (calculated from the dilution series) required to achieve maximal neutralisation. Each mAb clone and CDC combination group was repeated in triplicate experiments.

### Cross-linking mass spectrometry and data analysis

A 1:1 molar ratio of CDC antigen and mAb clone (either full-length antibody or Fab domain) was incubated in a Thermomixer at 37 °C with agitation at 500 rpm for 60 minutes. Subsequently, 1 mM DSS-H12/D12 or DSG-H6/D6 cross-linker was added to the samples, which were then incubated for 75 minutes at 37 °C and 1000 rpm. To quench the cross-linking reaction, 1 M ammonium bicarbonate was added. Thereafter, reduction and alkylation were performed by adding 8 M urea, 5 mM TCEP, and 10 mM iodoacetamide (IAA) as previously described. For protein digestion, 1 µg of lysyl-endopeptidase (FUJIFILM Wako) was initially added to the sample for pre-digestion at 37 °C and 500 rpm for 2 hours. The samples were then diluted with 100 mM ammonium bicarbonate and subjected to overnight digestion with 1 µg of trypsin at 37 °C and 500 rpm. The digestion was stopped by adding 40 µL of 10% formic acid. The resulting peptides were cleaned using C18 reverse-phase micro spin columns, dried in a SpeedVac for 4 hours, and stored at-20 °C until further use.

For LC-MS/MS analysis, approximately 1 µg of peptides from CDC-mAb cross-linked samples or 0.6 µg peptides from CDC-Fab cross-linked samples, as quantified by NanoDrop, were injected onto an Ultimate 3000 UPLC system connected to an Orbitrap Eclipse Tribrid mass spectrometer (Thermo Fisher Scientific). Each cross-linked sample was injected twice. Peptide separation was achieved using a linear gradient of Solvent A (0.1% formic acid) and Solvent B (0.1% formic acid, 80% acetonitrile), ranging from 5% to 25% B over 100 minutes at a constant flow rate of 300 nL/min. Datasets were collected using a data-dependent acquisition (DDA) method. Each cycle began with an MS1 scan over a range of 400-1,800 m/z at a resolution of 120,000, with standard AGC settings and auto-mode maximum injection time. This was followed by MS2 scans acquired in a 3 s cycle time, with a resolution of 30,000, standard AGC target, a maximum injection time of 100 ms, and a stepped normalised collision energy (NCE) of 21, 26, and 31. Only precursors with charge states between 3 and 8 were selected for fragmentation. Instrument performance was routinely monitored throughout the injection and analysis using a HeLa protein digest standard (Thermo Fisher Scientific).

The acquired cross-linking datasets were analysed using pLink ^59^ and MaxLynx ^60^ software. The cross-link search parameters were configured based on the specific linker modification types used in the experiment. XL peptide pair searching was conducted against a database containing either only the sequences of the target mAb and CDC antigen or an expanded database including common contaminant proteins. The search allowed for a maximum of two missed cleavage sites. Carbamidomethylation of cysteine residues was set as a fixed modification, while protein N-terminal acetylation and methionine oxidation were considered as variable modifications. For visualisation, xiVIEW ^61^ was employed to display inter-protein cross-links identified within the antigen-antibody (or Fab) complexes. The identified cross-linked sites were subsequently highlighted onto the three-dimensional structure of the corresponding CDC antigen using ChimeraX ^62^. The representative cross-linked spectrum match from each PLY-mAb pair was exported as peak list, and visualised and annotated accordingly in xiSPEC v2 ^63^.

### Hydrogen/deuterium exchange mass spectrometry and data analysis

HDX-MS experiments were conducted on either PLY or PFO antigen in both its apo (unbound) state and in complex state with different Fab fragments. To prepare the Fab-bound complex, PLY/PFO and various Fab were mixed in a 1:2 molar ratio in PBS to ensure epitope saturation. The samples were subsequently deuterated at 4 °C using a D_2_O-based HDX labelling buffer (Thermo Fisher Scientific) over five time intervals: 0, 30, 300, 3,000, and 9,000 seconds. For each labelling interval, the complexed (Fab-bound) PLY/PFO state was analysed in triplicate within a single continuous run, while the apo PLY state was repeated a total of six times (apo PFO was repeated three times) during the run. To map the paratope of 6E5 interacting to PLY antigen, the apo state and complex state were reversed. We then saturated the paratope with excessive PLY antigen. Other experimental setups followed the same protocol.

To quench the deuteriation, the samples were mixed with a quench buffer composed of 1% trifluoroacetic acid (Thermo Fisher Scientific), 0.2 M TCEP, and 4 M urea, and kept at 4 °C. The quenched samples were then injected at 4 °C into an on-line digestion using mixed protease column of Nepenthesin-2 and pepsin, and sample trapping system operating at a flow rate of 50 µL/min with 0.1% formic acid for 4 minutes. The digested peptides were trapped and cleaned on a PepMap 300 C18 trap column (Thermo Fisher Scientific) before being directed in-line to a reverse-phase analytical column. Chromatographic separation was performed at 1 °C using a mobile phase gradient that increased from 5% to 50% solvent B (0.1% formic acid, 95% acetonitrile) over 8 minutes, followed by an increase from 50% to 90% over the next 5 minutes. Eluting peptides were analysed on a Q Exactive Plus mass spectrometer (Thermo Fisher Scientific). Full MS scans were acquired at a resolution of 70,000 with an AGC target of 3e6 and a maximum injection time of 200 ms, covering an m/z range of 300 to 2,000. In parallel, peptide identification was performed by analysing an un-deuterated, digested PLY/PFO/6E5_Fab samples using DDA tandem MS. A peptide database, including sequence, charge state, and retention time information, was constructed with PEAKS Studio X (Bioinformatics Solutions Inc.).

HDX-MS data processing was carried out using HDExaminer v.3.4.2 (Sierra Analytics Inc.). For each identified peptide, individual charge states were considered for comparative analysis between the corresponding complex and the apo state. Deuterium incorporation levels were determined by measuring the increase in mass relative to the un-deuterated control peptide, using the most extensive deuteration (>9,000 s) as a reference. Manual inspection of spectra allowed the removal of peptides or experimental replicates with low scores, clear outliers, or inconsistent retention times. The curated data matrix was further analysed using Deuteros 2.0 ^64^ software to perform significance testing, differential analysis, and to generate various plots illustrating peptide kinetics and redundancy. Finally, ChimeraX was utilised to map and visualise the regions of PLY/PFO protected upon a given Fab binding, presenting the MS-derived structural epitope on the three-dimensional space of the corresponding antigen molecule. The same visualisation was applied to the HDX-MS-determined paratope peptide on the modelled 6E5 Fab domain.

### Multimodal data-driven HADDOCK analysis

An initial benchmark test was performed to evaluate the robustness of the data-driven antigen-antibody HADDOCK docking protocol. We used an external dataset in which the co-crystal structure had been solved (PDB: 6P3R), coupled with available HDX-MS data ^43^. For the input data, the antigen crystal structure was combined with either the extracted Fv crystal structure or an Fv model generated by ABodyBuilder2 ^65^. Epitope peptides identified by HDX-MS were specified as residues involved in the binding interface, together with the pre-defined 6 CDR hypervariable loops of the cognate Fv, with a requirement for the selected residue restraint that the percentage of the relative solvent accessibility (RSA) was no less than 15%. For the sampling parameters, 10,000 structures were generated for the initial rigid-body docking, followed by 500 models for sequential semi-flexible refinement, final refinement, and analysis, respectively. The top-ranked pairwise complex model was selected and compared with the co-crystal Fv-Ag structure. Complex models were superposed on the Fv structure to calculate antigen L-RMSD, and separately superposed on the interface to calculate i-RMSD in ChimeraX. Interface residues from both the input data combinations and the co-crystal structure were computed in PRODIGY ^66^ and highlighted with sidechain showing in ChimeraX.

Before modelling the mAb 6E5 Fab-PLY pairwise complex, two most prevalent cross-links were used to infer the epitope on PLY using the DisVis interaction analysis mode ^67^. Residues from both chains of the 6E5 Fab domain were renumbered to facilitate analysis. Briefly, DSG cross-links (PLY_Lys442-6E5_heavychain_Lys76 and PLY_Lys424-6E5_heavychain_Lys87) were applied with a Cα-Cα distance range of 0 to 24 Å. The inferred interface residue of PLY, referred to as a loosely defined epitope, was determined based on an interaction fraction greater than 0.5 among candidate accessible residues on PLY with an RSA, calculated from the PLY crystal structure in PyMOL, no less than 40%. For the input data, the antigen PLY crystal structure and the top-predicted AlphaFold-multimer ^38^ model of 6E5 Fab were used.

To explore different richness levels of restraint to guide docking, four scenarios of restraint combinations were defined as follows: i) pre-defined 6 CDR loops as the paratope with the loosely defined epitope derived from the preceding DisVis analysis; ii) HDX-MS determined paratope with the loosely defined epitope; iii) predefined 6 CDR loops as the paratope with the HDX-MS determined epitope; iv) paratope and epitope both determined by HDX-MS. The two DSG cross-links were imposed as the centre of mass distance constraints (Cα-Cα range of 0 to 24 Å) in all four scenarios throughout sampling and refinement, using the same parameters as in the benchmark analysis, to focus and maximise computational resources on the confined interaction space. The top-ranked 6E5 Fab-PLY model generated from each scenario was selected and visualised in ChimeraX, and interface characteristics and intermolecular contacts were computed with PRODIGY. For the case of mAb 6E5-PFO pairwise complex modelling, restraints equivalent to scenario iii were applied where DSG cross-link (PFO_Lys435-6E5_heavychain_Lys76, Cα-Cα range of 0 to 24 Å) was set as the centre of mass restraint, followed by the same analysis and visualisation procedure.

## Data Availability

The mass spectrometry proteomics data will be deposited to the ProteomeXchange Consortium via the PRIDE ^68^ partner repository upon completion of the peer-review process.

## Acknowledgements

We gratefully acknowledge the Swedish National Infrastructure for Biological Mass Spectrometry (BioMS), the SciLifeLab Integrated Structural Biology Platform for providing facilities and experimental support. We would also like to thank Dr. Christofer Karlsson, Dr. Anahita Bakochi for assistance.

---

**Extended Data Figure 1.**
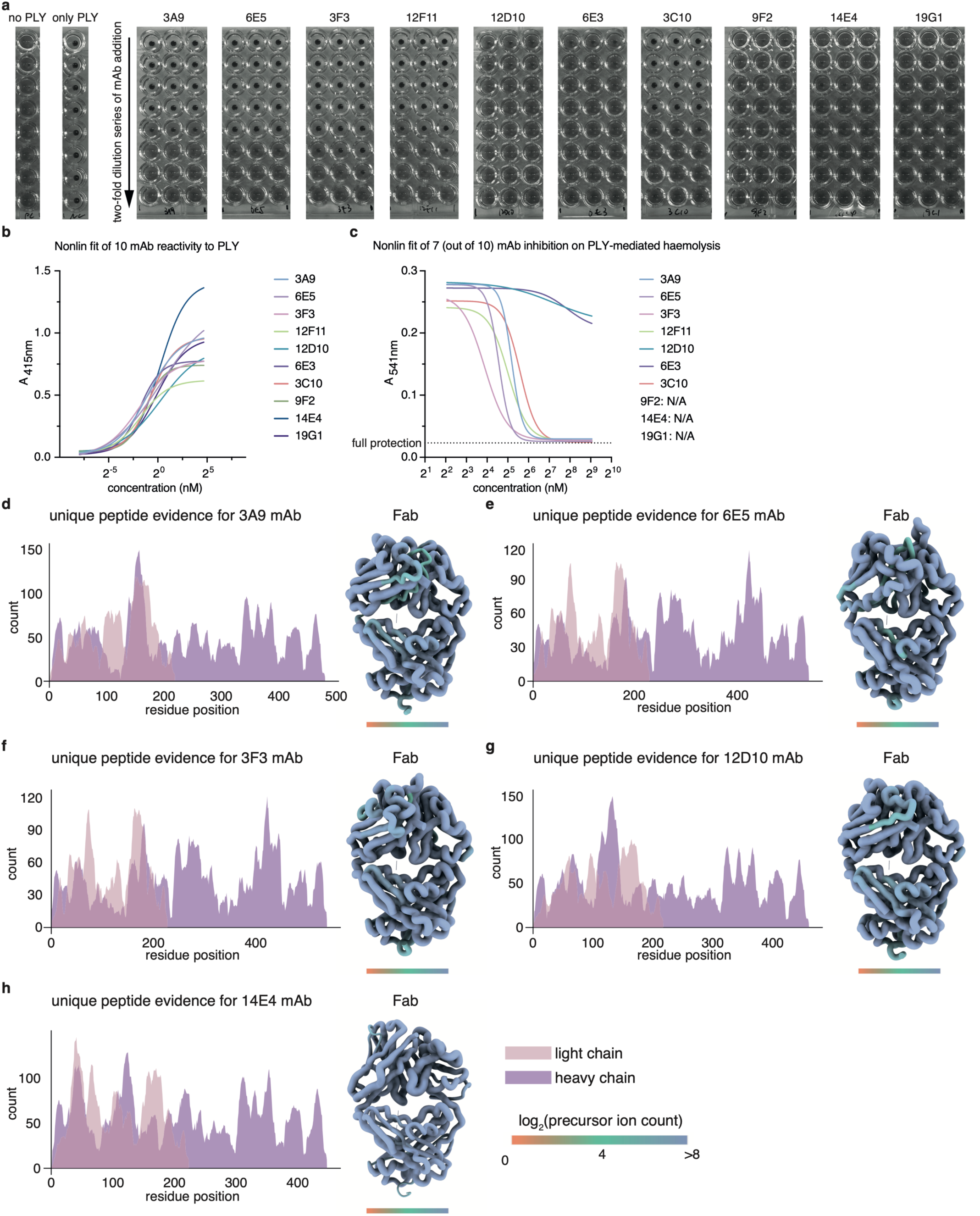
Binding specificity and haemolysis-inhibition potency of the PLY-mAbs and *de novo* MS sequencing result. **a)** Uncropped full monochromatic images showing pelleted, un-lysed RBCs at the bottom of the vials across the dilution series, representing the degree of protection, alongside the control groups on the left. **b)** Indirect ELISA with a 5-fold serial dilution was applied to calculate the EC50 values for the ten mAbs. A four-parameter non-linear regression analysis with variable slopes was applied for each mAb. **c)** Based on the absorbance measurements, the IC50 (half-maximal inhibitory concentration) of each protective mAb was determined from a two-fold serial dilution PLY-mediated haemolysis inhibition assay. The mAbs were subjected to multi-enzyme digestion using pepsin, trypsin, chymotrypsin, elastase, and alpha-lytic protease, followed by bottom-up mass spectrometry data-dependent acquisition. Candidate peptides were assembled using the Byonic SuperNova mAb sequencing workflow. The detected precursor ion support for each residue is plotted with the heavy and light chain position numbering overlaid. AlphaFold-Multimer was used to refine and build the hetero-dimeric Fab structure after modelling the variable regions with IgFold. Residues on the structure are coloured according to the supporting ion evidence. **d**-**h)** Five representative Fabs are shown here.

**Extended Data Figure 2.**
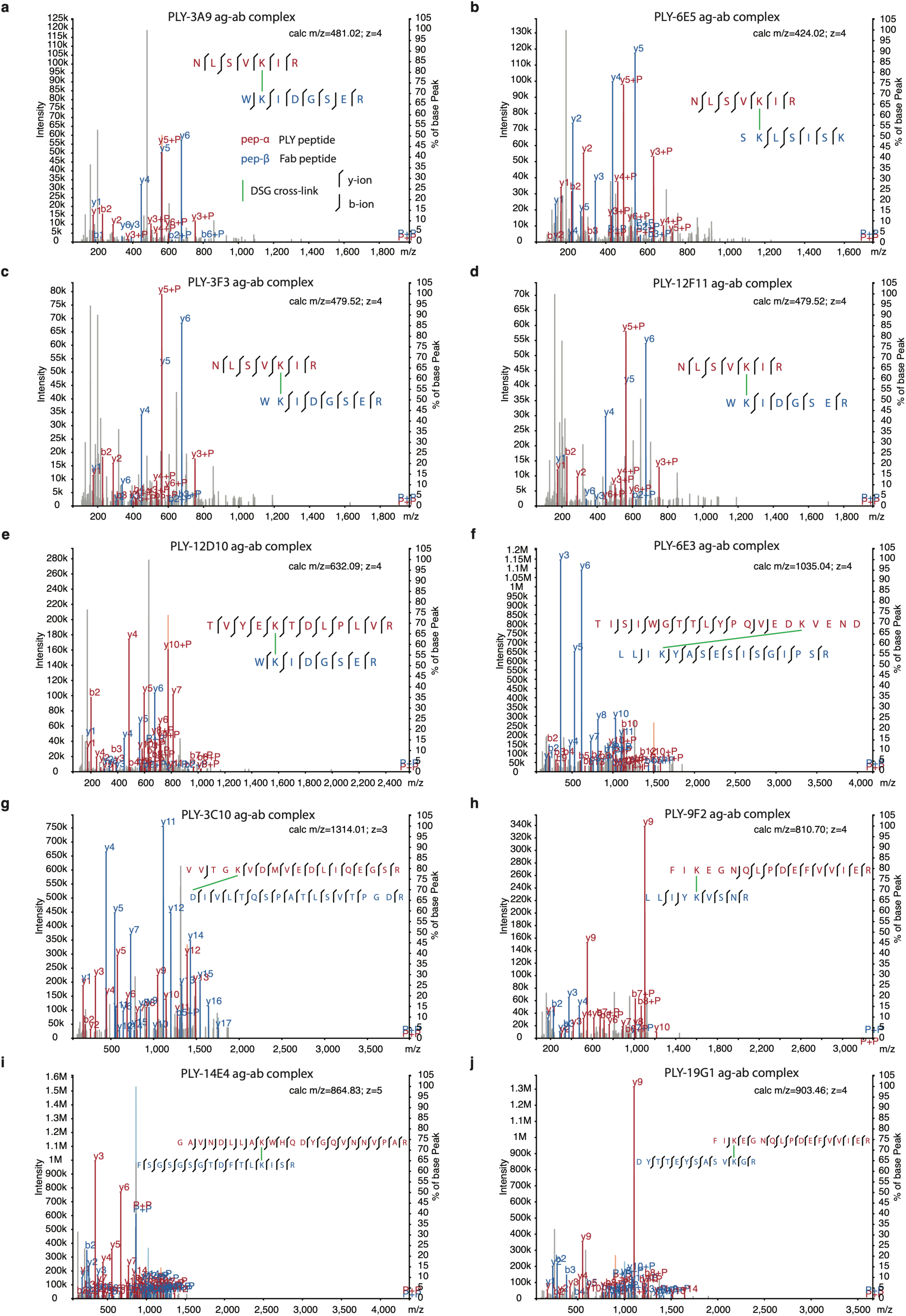
Annotated MS/MS spectra of DSG cross-linked peptide pairs found in the different PLY-mAb immune complexes. Representative MS/MS spectra for DSG cross-linked peptide pairs between PLY antigen and selected mAb clones. **a**) 3A9; **b**) 6E5; **c**) 3F3; **d**) 12F11; **e**) 12D10; **f**) 6E3; **g**) 3C10; **h**) 9F2; **i**) 14E4; **j**) 19G1. The matched fragmented ions are annotated and labelled according to the legend.

**Extended Data Figure 3.**
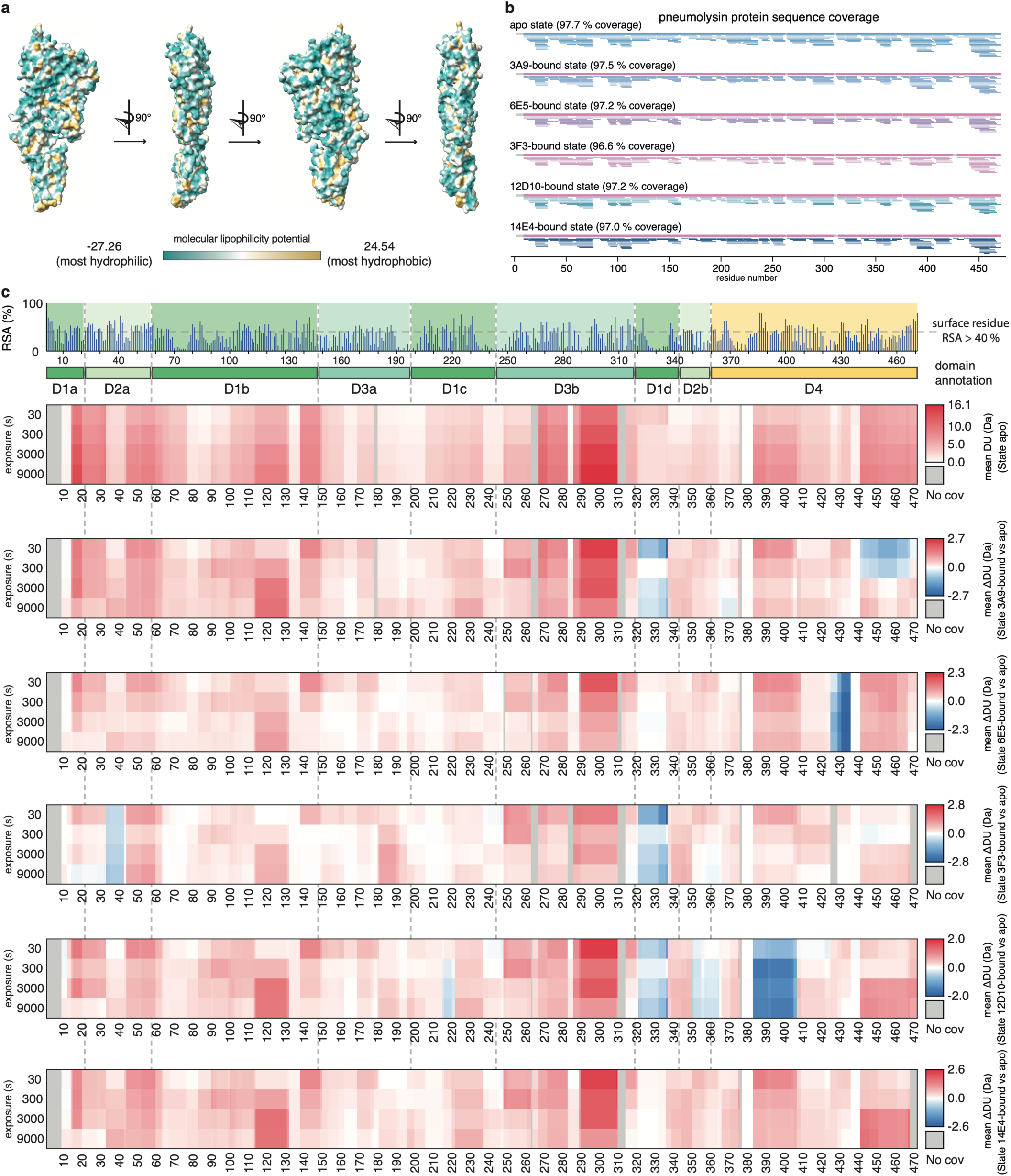
Hydrogen/deuterium exchange mass spectrometry analysis of pneumolysin protein during different Fab binding. **a)** The hydrophobicity (lipophilicity) of pneumolysin residues was calculated and visualised on a PLY surface representation from four different viewpoints. The gradient colour indicates the level of lipophilicity **b)** Coverage plots of the PLY protein sequence for all states from the HDX-MS experiment. Each bar represents a high-confidence peptide identification. **c)** Barcode plots illustrating the temporal dynamics of the mean deuterium uptake for the apo-state PLY and the mean differential deuterium uptake in the Fab-bound states for PLY in complex with 3A9, 6E5, 3F3, 12D10, or 14E4. Predicted relative solvent accessibility (RSA) is shown at top of the panel, aligned with domain annotation for PLY beneath.

**Extended Data Figure 4.**
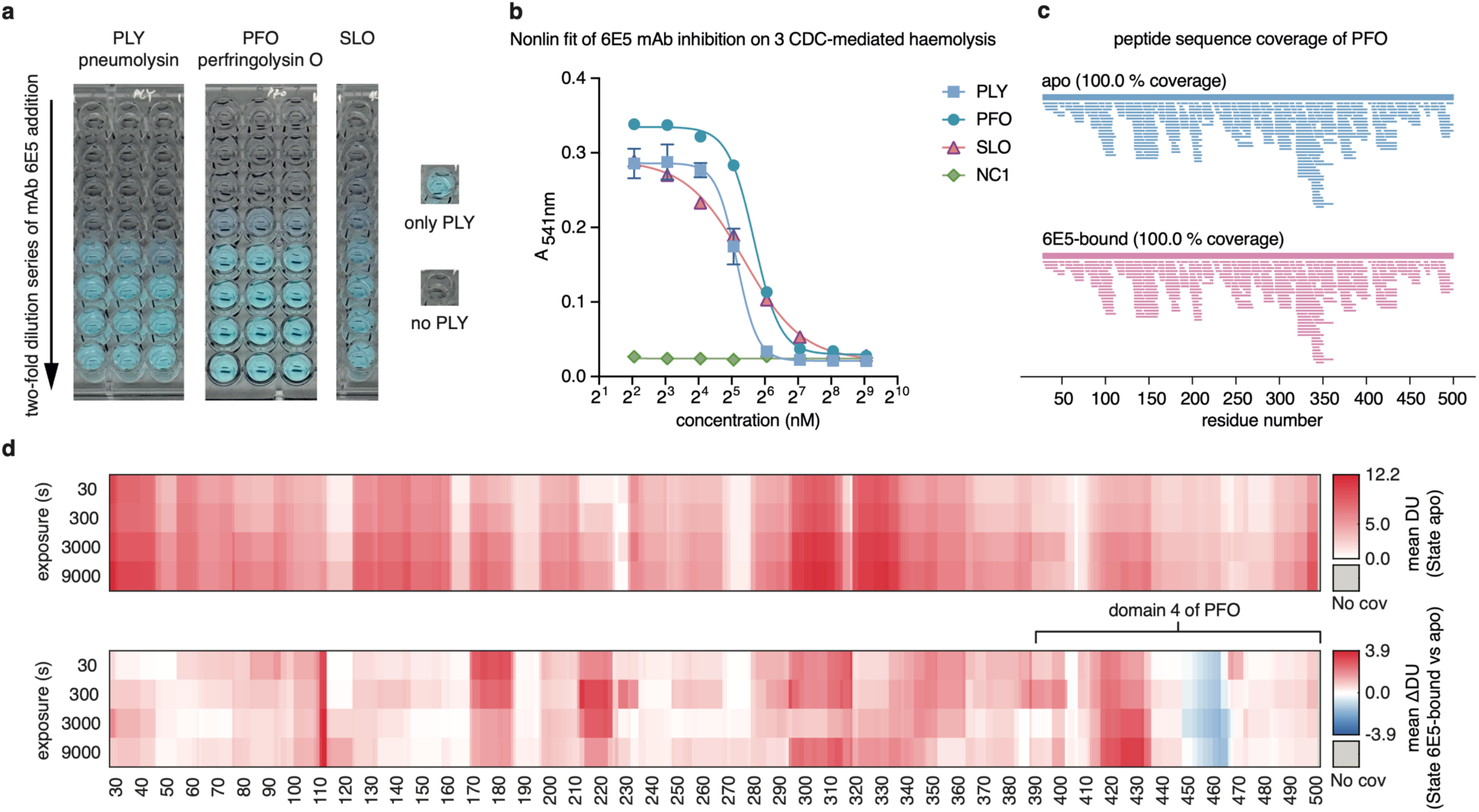
Cross-species CDC neutralising potency of mAb 6E5 and the corresponding cognate epitope peptide on PFO mapped by HDX-MS. **a)** Colour-inverted images showing transferred supernatant after centrifugation of un-lysed RBCs, reflecting the degree of protection, with the **b**) calculated IC50 (half-maximal inhibitory concentration) of mAb 6E5 on the right, determined from a two-fold serial dilution haemolysis inhibition assay using either PLY, PFO, or SLO. **c)** Coverage plots of the PFO, from *Clostridium perfringens*, across both apo and 6E5 Fab-bound states analysed by HDX-MS. Each bar represents a peptide identification. **d)** This panel of barcode plots illustrates the temporal dynamics of the mean deuterium uptake of apo state of PFO and the mean differential deuterium uptake in complex state of 6E5- bound PFO. Domain 4 of PFO spanning residue 391-500 is shown on the protein sequence.

**Extended Data Figure 5.**
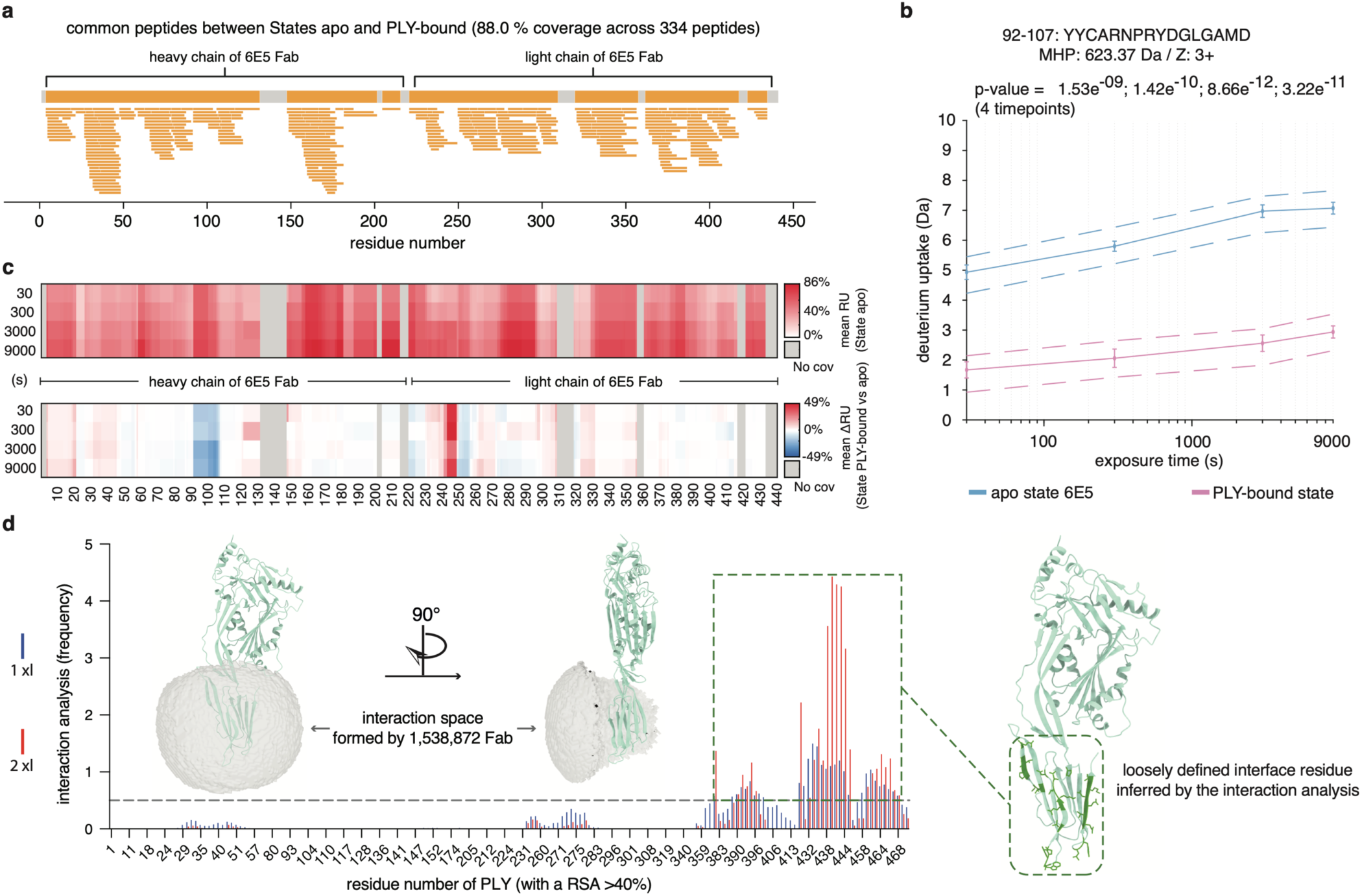
HDX-MS paratope mapping of mAb 6E5 in complex with PLY antigen and DisVis interaction analysis. **a)** A sequence coverage map of 334 common 6E5 peptides identified in both states of apo and the PLY-bound in HDX-MS. Each bar represents a peptide identification and absence of peptide coverage region is coloured in grey. **b)** The kinetic plot for one of the protected peptides in the 6E5 heavy chain variable region, showing deuterium uptake across four labelling intervals. Data are colour-coded for different states, with adjusted p-values annotated. MHP stands for the theoretical molecular weight of the peptide ion, and Z indicates the charge state. **c)** The top panel barcode plot displays the mean relative deuterium uptake (%) of 6E5 residues in apo state, while the bottom panel presents the mean differential deuterium change (%) upon PLY binding relative to the apo state. The colour gradient indicates the magnitude of deuterium change as a function of labelling interval. **d)** Here, two viewpoints of the PLY antigen with the potential interaction space formed by 1,538,872 6E5-PLY pair complexes are displayed, consistent with the two cross-link distance constraints. The bar plot shows the interaction frequency of each surface accessible residue on PLY from the DisVis interaction analysis, grouped by the number of cross-link distance constraints applied. The residues with a frequency index higher than 0.5 were extracted and designated as the inferred interface residues for downstream data-driven modelling.

**Supplementary Data Figure 1.**
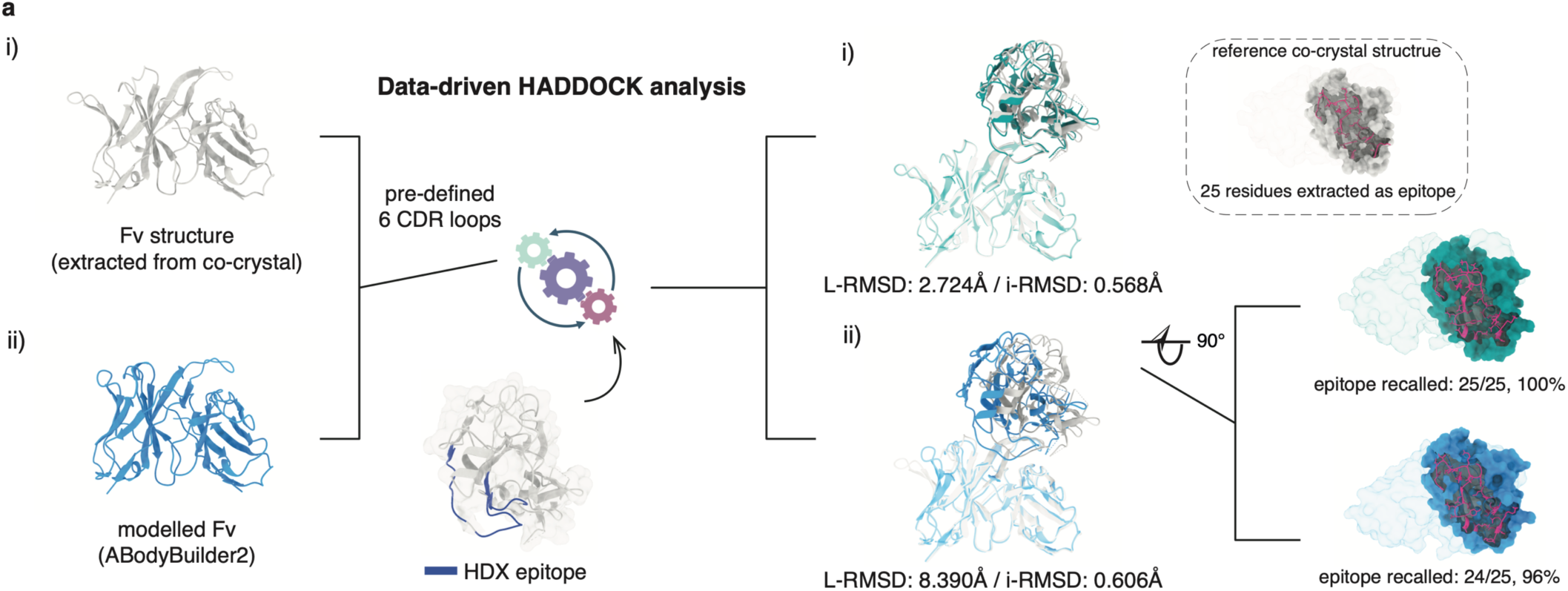
HADDOCK analysis benchmarked by an external dataset. **a)** Data-driven HADDOCK analysis was benchmarked using an external dataset for which the co-crystal structure of Fv-Ag pair complex is solved, shown in grey as the reference. Two docking runs were executed that differed solely in the Fv input structure: i) extracted directly from the co-crystal structure and ii) predicted from sequence, mirroring most real-world scenarios. Only HDX-MS-determined epitope peptide was used as the distance constraints for both runs. L-RMSD (ligand root-mean-square-deviation) and i-RMSD (interface root-mean-square-deviation) were calculated by superposing the corresponding top-ranked model onto the co-crystal structure accordingly. Interface residues from the modelled Fv-antigen complexes, from the bottom-up view of the antigen, are shown with sidechains coloured in hot pink, while all other non-interface residues are displayed in surface representation.

**Extended Data Table 1.** Usage and pairing of heavy and light chain VJ murine germline genes determined for the ten mAb clones. IGHV: immunoglobulin heavy chain variable gene; IGHJ: immunoglobulin heavy chain joining gene; IGKV: immunoglobulin kappa chain variable gene; IGKJ: immunoglobulin kappa chain joining cluster.

| mAb clone | IGHV | IGHJ | IGKV | IGKJ |
| --- | --- | --- | --- | --- |
| 3A9 | 5-12-1 | 1 | 10-94 | 1 |
| 12F11 | 5-12-1 | 4 | 10-94 | 1 |
| 9F2 | 5-6 | 1 | 1-110 | 1 |
| 12D10 | 5-9 | 4 | 15-103 | 4 |
| 19G1 | 7-1 | 1 | 4-74 | 1 |
| 14E4 | 9-2-1 | 3 | 1-117 | 2 |
| 3C10 | 3-8 | 1 | 5-45 | 5 |
| 6E5 | 2-6 | 4 | 8-30 | 5 |
| 3F3 | 2-3 | 4 | 5-43 | 4 |
| 6E3 | 2-3 | 4 | 5-45 | 5 |

